# Cellular Traction and Morphological Adaptation to Cell-Scale Stiffness Gradients on Length-Encoded Micropillar Arrays

**DOI:** 10.64898/2026.10.22.760892

**Authors:** Prabuddha De Saram, Nam-Trung Nguyen, Sharda Yadav, Navid Kashaninejad

**Affiliations:** Queensland Quantum and Advanced Technologies Research Institute, Griffith University, 170 Kessels Road, Nathan, QLD 4111, Australia; School of Engineering and Built Environment, Griffith University, 170 Kessels Road, Nathan, QLD 4111, Australia

**Keywords:** Micropillar arrays, Cell traction force, Stiffness gradients, Durotaxis, Two-photon polymerisation (2PP), Mechanotransduction

## Abstract

In tumour microenvironments, cancer cells routinely encounter abrupt and spatially heterogeneous stiffness variations at the scale of individual cells. How cells sense and respond to such steep, local stiffness gradients is not well understood. Here, we developed length-encoded micropillar arrays in which stiffness is set by the effective pillar length, while the pillar tops remain at the same height and spacing. Pillars with diameters of 2, 2.5 and 3 µm gave stiffness values of 4.0–82.6 nN/µm (effective modulus 2.9–39.4 kPa), varying periodically over distances comparable to the size of individual cancer cells. This platform allows simultaneous measurement of cell morphology, alignment, spreading, pillar displacement, and traction forces. In MDA-MB-231 breast cancer cells, pillar displacement showed a biphasic response. It decreased with increasing stiffness up to about 10 nN/µm and then remained close to 0.5 µm. In contrast, traction forces increased across the full stiffness range. Cells contacted soft pillars more than stiff pillars, with about 23% greater coverage of soft regions. Cells also avoided large stiffness differences along their length: among cells whose two ends sat on pillars of equal stiffness, 48–50% lay along a single stiffness row, compared with about 14% expected by chance. Cell alignment was enriched along the two principal array directions (27–31% of cells within 10°) and increased slightly with pillar diameter. Length-encoded micropillar arrays therefore provide a platform for studying how cells respond to cell-scale mechanical heterogeneity, with relevance to cancer invasion.

## 1. Introduction

Cells continuously sense and respond to the mechanical properties of their surrounding environment, a process known as mechanotransduction. Substrate stiffness, viscoelasticity, topography, geometry, and external forces influence cellular responses, including proliferation and apoptosis, cell morphology, migration, and differentiation. Understanding how cells integrate these cues is essential for advancing biomaterials design, tissue engineering, and mechanobiology [1, 2].

Among various mechanical properties of the extracellular matrix (ECM), stiffness has emerged as a key parameter that directly affects cell migration, spreading, morphology, and differentiation through receptor-mediated mechanotransduction [3]. Mechanosensing primarily begins at integrin–ECM adhesions, where ligand binding transmits force across focal adhesions to the cytoskeleton and nucleus [4]. Following integrin binding to the ECM, actomyosin generates intracellular contractile forces. ECM resistance places tension on integrins and associated proteins, triggering molecular signalling through force-dependent conformational changes, such as protein unfolding. These mechanical signals propagate through the cytoskeleton and activate mechanobiological pathways that regulate cellular responses [4, 5]. This process is dynamic, as cells actively remodel the ECM through both biochemical activity and mechanical forces.

Stiffness-tuned hydrogel substrates, together with traction-force microscopy, have been widely used to study stiffness-based cell response [6-8]. While continuous substrates better mimic the continuous ECM, limitations include difficulty in creating precise, steep stiffness variations, controlling geometrical cues and measuring cellular forces [9]. Micropillar arrays provide a powerful tool for studying cell–substrate interactions. They enable direct quantification of cellular traction forces (CTFs) from pillar deflections and provide precise spatial control over substrate properties. Each micropillar serves as an independent force sensor, enabling high-resolution mapping of cell-induced forces. Individual micropillars can also be engineered to provide specific mechanical or geometrical cues to cells [9].

In vivo, cells frequently encounter stiffness gradients at soft–hard tissue interfaces, during embryogenesis and wound healing, and within the tumour microenvironment [10, 11]. Previous studies have shown that stiffness gradients regulate cell migration, elongation, and polarisation [12-14]. However, most studies investigate bulk or continuous stiffness gradients using hydrogels. The combined effects of discrete cell-scale stiffness modulation and topographical variation remain poorly understood [10, 15]. Furthermore, it remains unclear how localised mechanical heterogeneity regulates CTFs and morphological adaptation. Investigating these responses at the cell scale is important for understanding how cells integrate simultaneous mechanical and topographical cues within complex microenvironments. This knowledge may help develop improved biomimetic platforms for mechanobiology, tissue engineering, and disease modelling.

Micropillar stiffness is most often varied by changing pillar diameter. However, this also changes the focal adhesion area [16], which affects the traction force a cell exerts on each pillar. Varying the effective pillar length instead changes stiffness without changing the pillar diameter or the contact area. Length-varied pillars therefore allow stiffness sensing to be studied independently of adhesion geometry. However, controlling pillar length is difficult with conventional fabrication methods, and few studies have used it to regulate cell behaviour. Earlier work was limited to simple designs, such as a single step in pillar length [17]. Wang et al. produced a smooth stiffness gradient by transferring a capillary-formed height profile between micro-ridges to the micropillar base, and observed stiffness-dependent cell migration [18]. However, this approach offers limited control over the stiffness profile and cannot easily produce steep, cell-scale gradients. That study also focused only on migration over relatively large-scale stiffness variations.

In our previous work, we introduced a scalable fabrication strategy for height-varying microstructures and demonstrated their ability to tune surface wettability through controlled height gradients [19]. More recently, we applied height-varying micropillar arrays to passive micromixing and showed that asymmetric height profiles enhanced transverse transport and interfacial stretching under low-Reynolds-number conditions [20]. These studies demonstrate the versatility of height-varying microstructures for regulating interfacial and transport phenomena. However, their potential as cell-scale mechanobiology platforms for local stiffness sensing, morphological adaptation, and traction force generation remains largely unexplored.

Investigating these effects requires an adherent cell model that can actively interact with its mechanical environment through cytoskeletal reorganisation, spreading, migration, and traction force generation. MDA-MB-231 cells, a widely used human triple-negative breast cancer cell line with a highly migratory phenotype, provide a suitable model for examining such mechanically regulated cellular responses [21].

In this work, we investigate how MDA-MB-231 cells respond to micropillar substrates with repeating cell-scale stiffness gradients generated by varying effective pillar length. We examine cellular morphology, alignment, spreading behaviour, pillar displacement, and cell traction force generation as functions of pillar stiffness and diameter. We combine quantitative cell-morphology analysis with traction-force measurements derived from pillar deflection to determine how substrate mechanical heterogeneity influences cell morphology and force generation. Our platform also enables us to investigate cellular responses to steep, periodic stiffness variations over length scales comparable to individual cells.

## 2. Materials and Methods

Unless otherwise specified, all chemicals and reagents were obtained from Sigma-Aldrich, Australia, and used as received.

### 2.1. Micropillar Array Design

The design of the micropillar arrays primarily aims to modulate the substrate stiffness independently of other geometrical parameters perceived by cells (**Fig. 1**). The length was modulated in a periodic pattern to vary the effective substrate stiffness. The top surface of the pillars is maintained at the same level, while the base depth changes, so pillar length varies. Pillar length increases from 8 to 12 µm over five levels (8, 9, 10, 11 and 12 µm), and then decreases back to 8 µm in the same number of steps. This pattern repeats in one direction, whereas in the perpendicular direction, each pillar row maintains the same length.

**Fig. 1.**
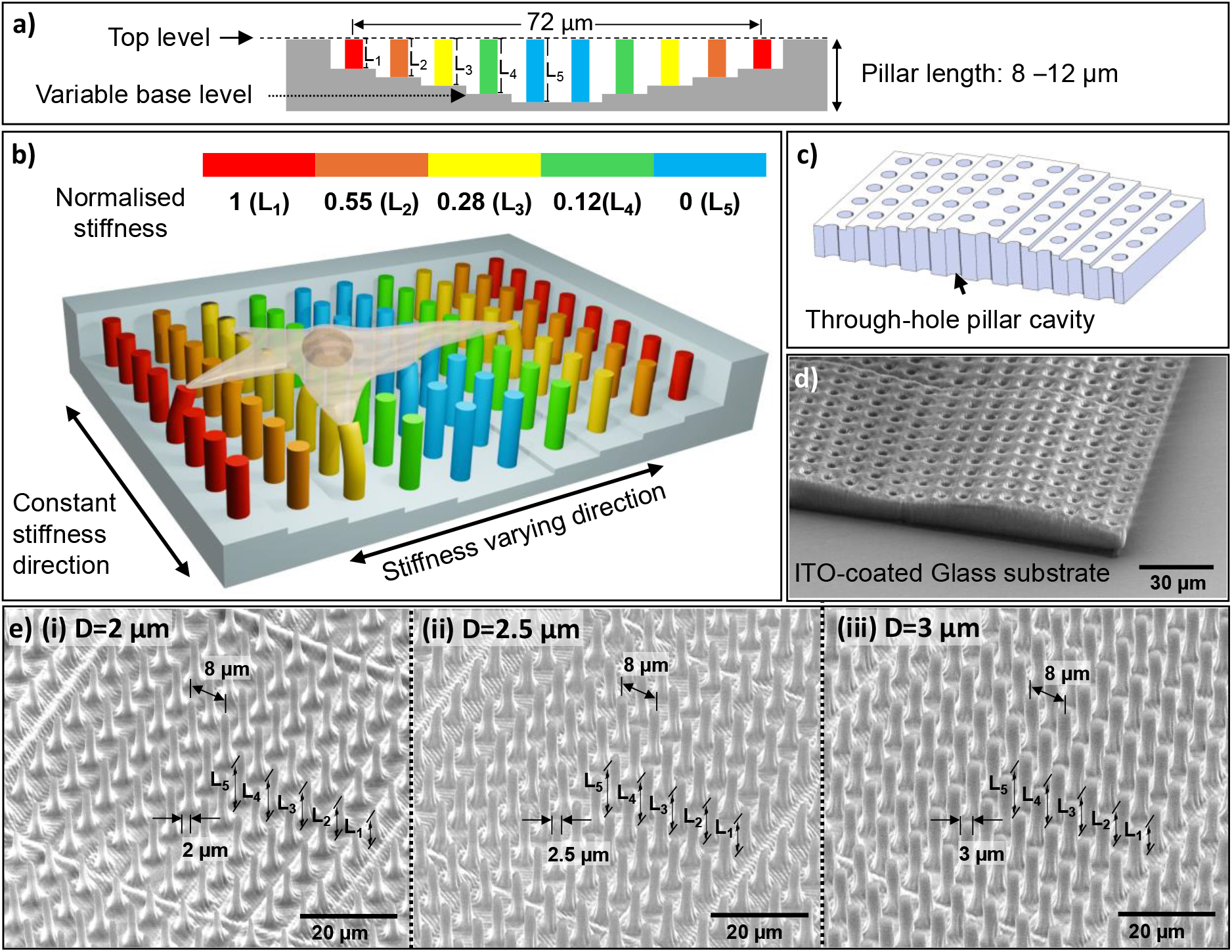
Micropillar array. (a) Schematic illustration of the micropillar array design. L_1_, L_2_, L_3_, L_4_, and L_5_ refer to the effective pillar lengths from the base to the top. (b) Schematic representation of the micropillar array parameter naming convention. The colour map shows the stiffness variation within the substrates. (c) Micropillar array mould design. Cylindrical through-hole cavities are fabricated in the IP-S resin, with the underlying ITO-coated glass substrate forming the base of each cavity, resulting in a smooth pillar top surface after release of the PDMS device. (d) Scanning electron microscopy (SEM) images of the two-photon lithography (2PP)-fabricated negative pillar mould. The moulds were surface-treated and subsequently used for direct PDMS casting. (e) SEM images of the fabricated PDMS micropillar arrays with pillar diameters of (i) 2 μm, (ii) 2.5 μm, and (iii) 3 μm.

The centre-to-centre distance between pillars is 8 µm. We used three substrates with corresponding pillar diameters of 2 µm, 2.5 µm, and 3 µm for the experiments. The pillar tops are aligned with the substrate surface, as the pillars are flush-integrated within a well-like structure (**Figs. 1a, 1b**). This design reduces the likelihood of pillar collapse under external shear forces applied to the substrate surface, such as during washing.

### 2.2. Micropillar Substrate Fabrication

We fabricated micropillars using poly(dimethylsiloxane) (PDMS) soft lithography with moulds created via two-photon polymerisation (2PP; Nanoscribe Photonic Professional GT2, Nanoscribe GmbH, Germany). Negative master moulds were fabricated using the Nanoscribe commercial resin (IP-S) on indium tin oxide (ITO)-coated glass substrates (**Figs. 1c,d**). The fabricated moulds were UV-cured for 1 minute to ensure complete polymerisation, followed by vacuum coating with trichloro(1H,1H,2H,2H-perfluorooctyl)silane (97%, Sigma-Aldrich, product no. 448931) to facilitate PDMS release after casting. The micropillar arrays were moulded from a 7:1 (w/w) mixture of PDMS base and curing agent and cured overnight at 60 °C.

### 2.3. Cell Culture

Before cell seeding, the substrates were treated with oxygen plasma for 15 s. Then micropillar tops were coated with 50 µg/mL human plasma fibronectin (Merck Life Science, Australia) by directly dispensing a droplet of solution onto the pillar array and incubated for 10 min at room temperature to support focal adhesion formation. Next, the substrates were immersed in 0.1% bovine serum albumin (BSA) for 1 h under vacuum at room temperature to ensure that the solution filled the gaps between the pillars. This blocked the pillar side walls and reduced non-specific cell adhesion. The coated substrates were placed in a 12-well plate, and MDA-MB-231 cells were seeded at 60,000 cells per well. Cells were cultured at 37 °C and 5% CO_2_. Traction forces were measured 1 day after seeding, and cell morphology was analysed 2 days after seeding.

### 2.4. Imaging and Microscopy

Live cells were imaged by brightfield (BF) microscopy (Olympus IX73, Olympus Corporation, Tokyo, Japan). Lower-magnification (20× objective) images were used for cell morphology analysis, and higher-magnification (60× objective) images were used to measure pillar deflection.

For scanning electron microscopy (SEM), cells were fixed with 2.5% glutaraldehyde and dehydrated through a graded ethanol series. Micropillar arrays and fixed cells were imaged with an Apreo 2 SEM (Thermo Fisher Scientific, Waltham, MA, USA) in low-vacuum mode. SEM images were used to measure pillar geometry and to confirm cell attachment to the pillar tops (**Fig. 2a**). SEM images of cells were not used for quantitative displacement analysis.

**Fig. 2.**
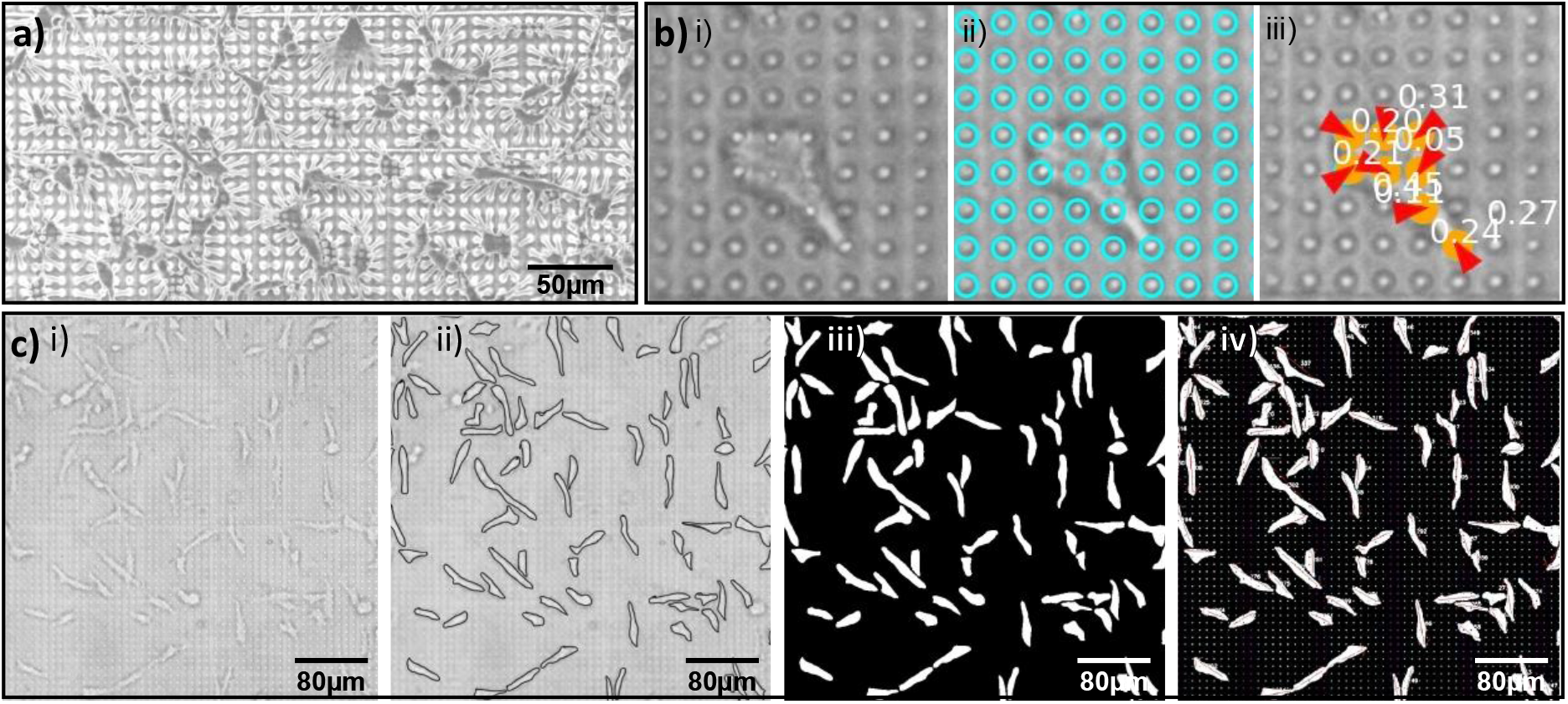
Processing of cell and pillar images. (a) SEM images of cells attached to micropillar arrays. Cell shrinkage during sample dehydration and drying resulted in exaggerated pillar deformations. These images were used solely to confirm cell attachment to the pillar tops and were not used for quantitative displacement analysis. (b) Representative image-analysis workflow used for pillar displacement detection and traction force calculation: (i) brightfield image of cells on pillars, (ii) ideal grid overlay generated from free pillars, and (iii) measured pillar displacement relative to the ideal grid. (c) Cell morphology analysis procedure. Cell outlines were identified from brightfield images (i, ii), and binary masks (iii) were generated using ImageJ. The binary masks, together with the corresponding pillar coordinates, were subsequently analysed using a custom Python script to quantify cell morphological parameters. In panel (iv), the small dots represent individual pillars, with colour indicating the corresponding pillar stiffness.

### 2.5. Pillar Deflection and Traction Force Analysis

Traction forces were measured from the higher-magnification BF images. Pillar tops appear as bright spots. An ideal grid was fitted to undeflected pillars, and the displacement of each pillar top was measured relative to this grid. The mean centre-to-centre pillar spacing (designed as 8 µm) was used to calibrate the spatial scale for displacement measurements (**Fig. 2b**).

From the measured pillar deflection δ, the traction force F was calculated as:

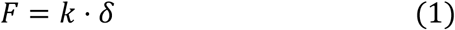

where *k* is the stiffness of the pillars. Pillar stiffness depends on the pillar length (*L*), the second moment of area (*I*), and the Young’s modulus (*E*) of the pillar material. The stiffness was calculated based on Euler–Bernoulli beam theory. SEM images (**Fig. 1e**) revealed a tapered base region formed during the 2PP fabrication process. Nominal pillar diameter, base diameter and taper angle were measured from the SEM images and incorporated into the calculation of stiffness using a variable cross-sectional beam model (refer to Section S1 of the Supplementary Information).

Additionally, substrate warping-induced tilting at the pillar base was considered to assess its potential influence on the CTF calculation [22]. However, the enlarged pillar–substrate contact region produced by the tapered base substantially reduces the contribution of substrate warping. For example, for a tapered-base pillar with 2-µm diameter and 10-µm length, the calculated stiffness was approximately 7.37 nN/µm. Incorporating the estimated substrate-warping contribution reduced the stiffness to approximately 7.26 nN/µm, corresponding to a difference of only about 1.5%. Across the full range of pillar geometries used in this study, the largest estimated reduction in stiffness due to substrate warping was approximately 6%, representing the possible maximum expected influence of this effect. Therefore, the contribution of substrate warping was considered relatively minor and was not included in the calculation of final stiffness.

The Young’s modulus of the 7:1 PDMS was measured by uniaxial tensile testing of a specimen cast in a 3D-printed mould. Force and elongation were recorded with a commercial delaminator test system (DTS Company, Menlo Park, CA, USA), modified in-house for tensile testing, comprising a linear actuator and a force sensor (Section S1 of the Supplementary Information).

Calculated traction force vectors were grouped for individual cells or cell clusters. To reduce residual global measurement errors arising from image registration or uncertainty of grid-fitting, a force-balance correction was applied so that the net vector sum of forces within each isolated cell or cell cluster approaches zero.

The effective Young’s modulus (*E*_*eff*_) of a continuous substrate equivalent to the pillar array can be estimated as: [23-25]

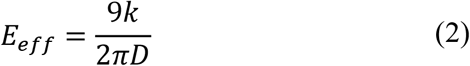

where *D* is the pillar diameter and *k* is the individual pillar stiffness calculated above. This equivalent modulus enables comparison of the mechanical properties of micropillar arrays with those of continuous gel substrates and natural extracellular matrices.

We define the normalised stiffness *k*^*\**^ to describe the relative stiffness of each pillar length within a substrate:

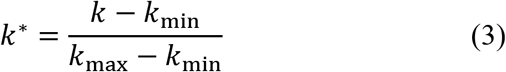

where *k*_min_ and *k*_max_ are the respective minimum and maximum pillar stiffness of a given substrate. Thus, *k*^*\**^ ranges from 0 for the softest pillar to 1 for the stiffest pillar. Although the absolute pillar stiffness differs between substrates due to the different diameters, the identical pillar-length pattern used across all substrates results in the same *k*^*\**^ values for corresponding pillar lengths. The relative stiffness variation experienced across an elongated cell is,

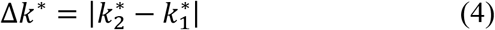

where 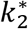 and 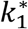 are the respective normalised stiffness values at the two ends of an elongated cell. The absolute value of the normalised stiffness difference was used to avoid negative values. Accordingly, Δ*k*^*\**^ = 0 represents no stiffness difference across the cell, whereas larger Δ*k*^*\**^ values indicate a greater relative stiffness variation.

### 2.6. Cell Morphology Analysis

Cell morphology was analysed using brightfield images capturing the full micropillar array area. A binary mask of cells was generated using Fiji [26] (ImageJ) and manually validated against images at different focal planes to ensure accurate cell identification; any misidentified regions were corrected manually (**Fig. 2c**).

We analysed the processed binary masks using custom Python scripts to extract morphological parameters, including cell spread area, aspect ratio, circularity, and orientation angle, based on ellipse fitting of individual cell boundaries. In addition, we incorporated a pillar grid with stiffness encoding into the analysis to map cell regions to corresponding substrate stiffness. This enabled quantification of cell spreading and morphological variation as a function of local pillar stiffness. Cell alignment was quantified as the angular deviation between the major axis of each fitted ellipse and the stiffness-varying direction (SVD).

### 2.7. Statistical Analysis

We analysed the data with custom Python scripts. Results are presented as mean ± standard deviation (SD) or mean ± standard error of the mean, as indicated in the figure captions. Statistical significance was evaluated using t-tests, with p < 0.05 considered statistically significant. Three independent datasets were analysed for each condition (*n* = 3). For each replicate, parameters were calculated independently and subsequently pooled for combined statistical analysis.

## 3. Results and Discussion

In this study, we examined cell morphological changes and cellular traction force patterns in response to the change in pillar stiffness. The SVD and constant-stiffness direction (CSD) refer to the two orthogonal directions as indicated in **Fig. 1b**.

### 3.1. Traction Force Variations

Assuming small tip displacement and a tapered base, we estimated pillar stiffness numerically by integrating the Euler–Bernoulli beam theory equation over the pillar length using a custom Python script (see Section S1 of the Supplementary Information). Table 1 summarises the calculated pillar stiffness. For the 7:1 PDMS formulation used to fabricate the micropillar arrays, the Young’s modulus was experimentally determined to be 2.2 MPa. The corresponding effective Young’s modulus (*E*_eff_) values of the micropillar arrays fall within the biologically relevant ECM stiffness range reported for breast cancer cells [27, 28].

**Table 1.**
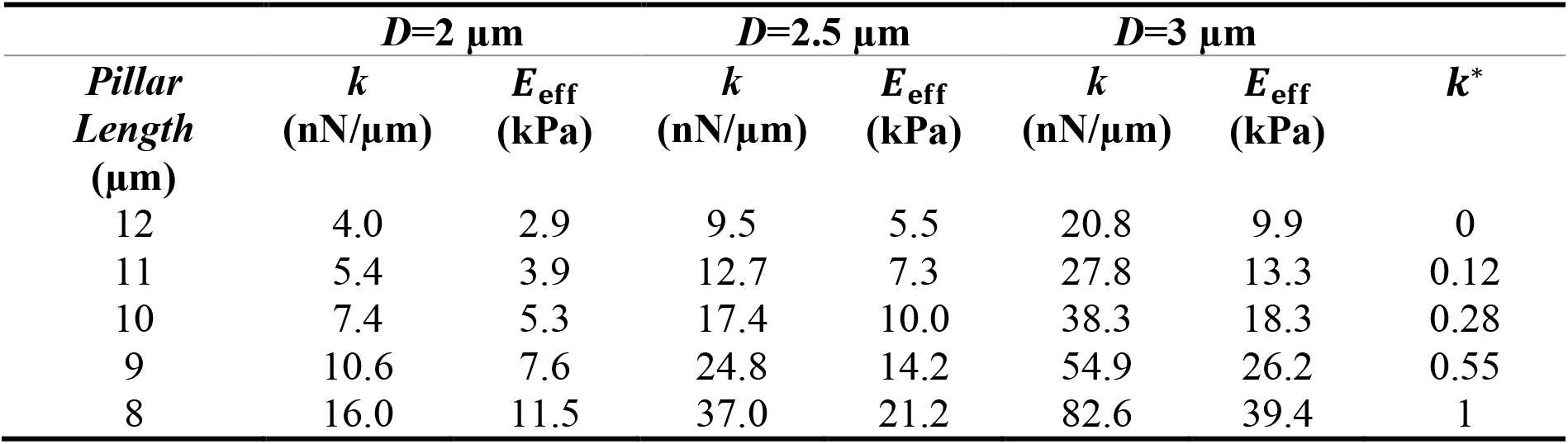
Calculated individual pillar stiffness (*k*), effective Young’s modulus of a continuous substrate (*E*_eff_) and normalised stiffness (***k***^*\**^) for pillar lengths

**Fig. 3a** depicts displacements caused by cells across different pillar stiffness values and diameters. The results reveal a biphasic displacement response, with a transition occurring around ∼10 nN/µm. The 2-µm pillars fall predominantly within the first phase, whereas the 2.5- µm and 3-µm pillars lie mainly in the second phase. In the initial phase, pillar displacement decreases with increasing stiffness, although at a progressively reduced rate. Beyond the transition, displacement reaches an approximately constant value of ∼0.5 µm and is maintained across higher stiffness values.

**Fig. 3.**
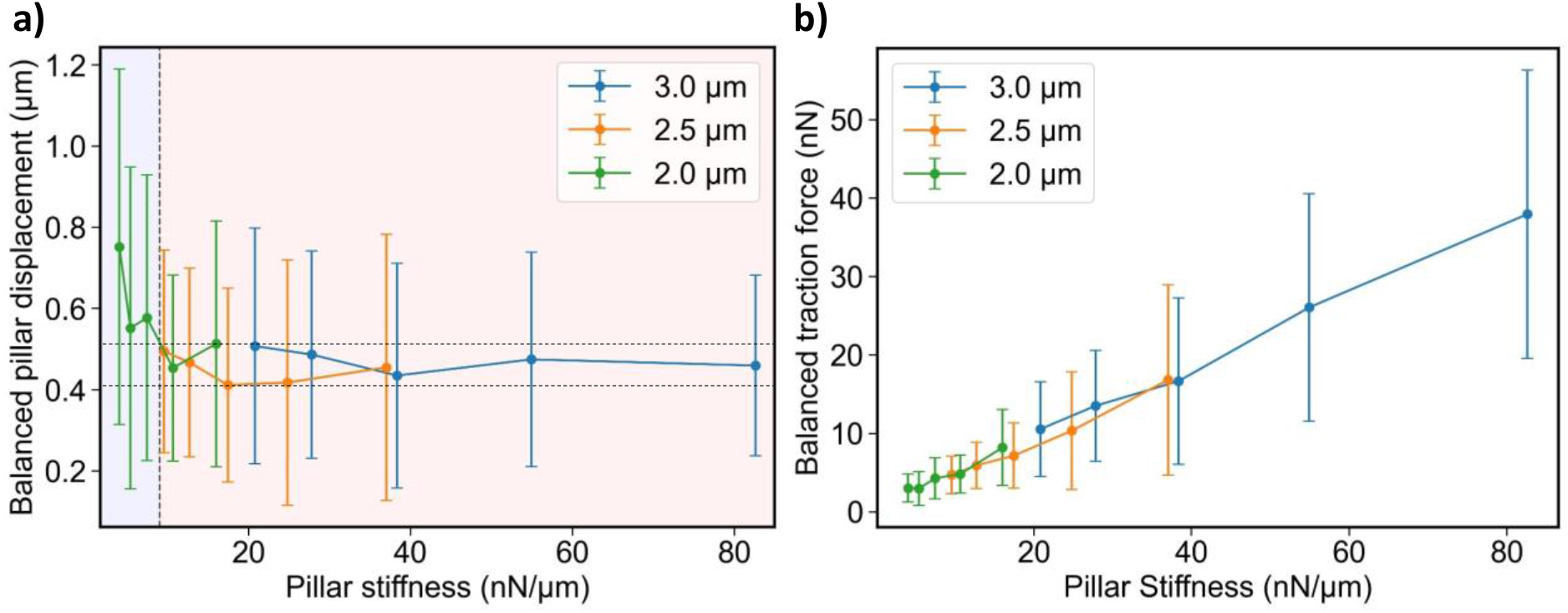
(a) Cell-induced pillar displacement as a function of pillar stiffness for arrays with different pillar diameters. (b) Cellular traction force (CTF) as a function of pillar stiffness for the corresponding micropillar arrays. Error bars represent the standard deviation of the measurements.

The measured pillar-top displacement was converted into CTF using the stiffness values listed in Table 1. Consistent with previous studies, CTF shows strong sensitivity to substrate stiffness, even when cells span multiple stiffness values simultaneously [29]. **Fig. 3b** presents the variation of balanced CTF with pillar stiffness. The results show a monotonic increase in traction force with increasing stiffness, with a gradual rise at low stiffness, a more pronounced increase at intermediate stiffness, and a slightly reduced rate of increase at higher stiffness, suggesting a tendency toward force saturation.

Previous studies have reported that cells tend to maintain a relatively constant level of substrate deformation across varying stiffness conditions [16, 24]. The displacement plateau observed in the second phase supports this concept, indicating that cells may reach a mechanoregulated state in which traction forces are actively adjusted to maintain a preferred level of deformation (∼0.5 µm) while supporting stable focal adhesions. Notably, the average displacement level remains within a similar range despite differences in pillar diameter between 2.5 µm and 3 µm. Given that pillar stiffness scales with the fourth power of the diameter, while focal adhesion area scales approximately with the square of the diameter, this behaviour highlights the role of active mechanosensing and feedback regulation in maintaining consistent deformation.

In contrast, the first phase represents a regime in which pillar displacement decreases with increasing stiffness, while the corresponding CTFs increase at a relatively slower rate. Focal adhesions require sufficient mechanical resistance from the extracellular environment to fully develop and mature [30]. In this regime, softer pillars may undergo larger passive deformation under CTFs while providing insufficient mechanical resistance to support mature focal adhesion formation and transition to the mechanoregulated state. Furthermore, less organised focal adhesions may contribute to the relatively large standard deviation in pillar displacement observed at the first few stiffness values.

### 3.2. Spreading Patterns on the Substrates

These substrates have steep stiffness gradients with short spatial periods. As a result, cells do not experience a consistent mechanical cue over long distances to guide migration. The wavelength of stiffness variation is comparable to the size of an elongated cell. As a result, even if a cell migrates, it cannot fully avoid or reach a preferred stiffness region. This leads to distinctive cell spreading behaviour rather than directed migration. Accordingly, no clear cell accumulation was observed on the substrates.

To quantify the spreading behaviour, we analysed the distribution of pillars covered by cells across all normalised stiffness values (**Fig. 4a**). On average, 44.7% of pillar tops were covered in the softer regions (11 and 12 µm pillars), compared with 36.5% in the stiffer regions (8 and 9 µm pillars), representing a ∼23% higher coverage in the softer regions. This trend shows that across all substrates, cells tend to maintain more contact with softer pillars. In contrast, numerous studies have shown that cells cultured on deformable substrates preferentially migrate towards stiffer regions in response to substrate stiffness gradients, a phenomenon known as positive durotaxis. This behaviour has also been reported for several cancer cell types [31, 32].

**Fig. 4.**
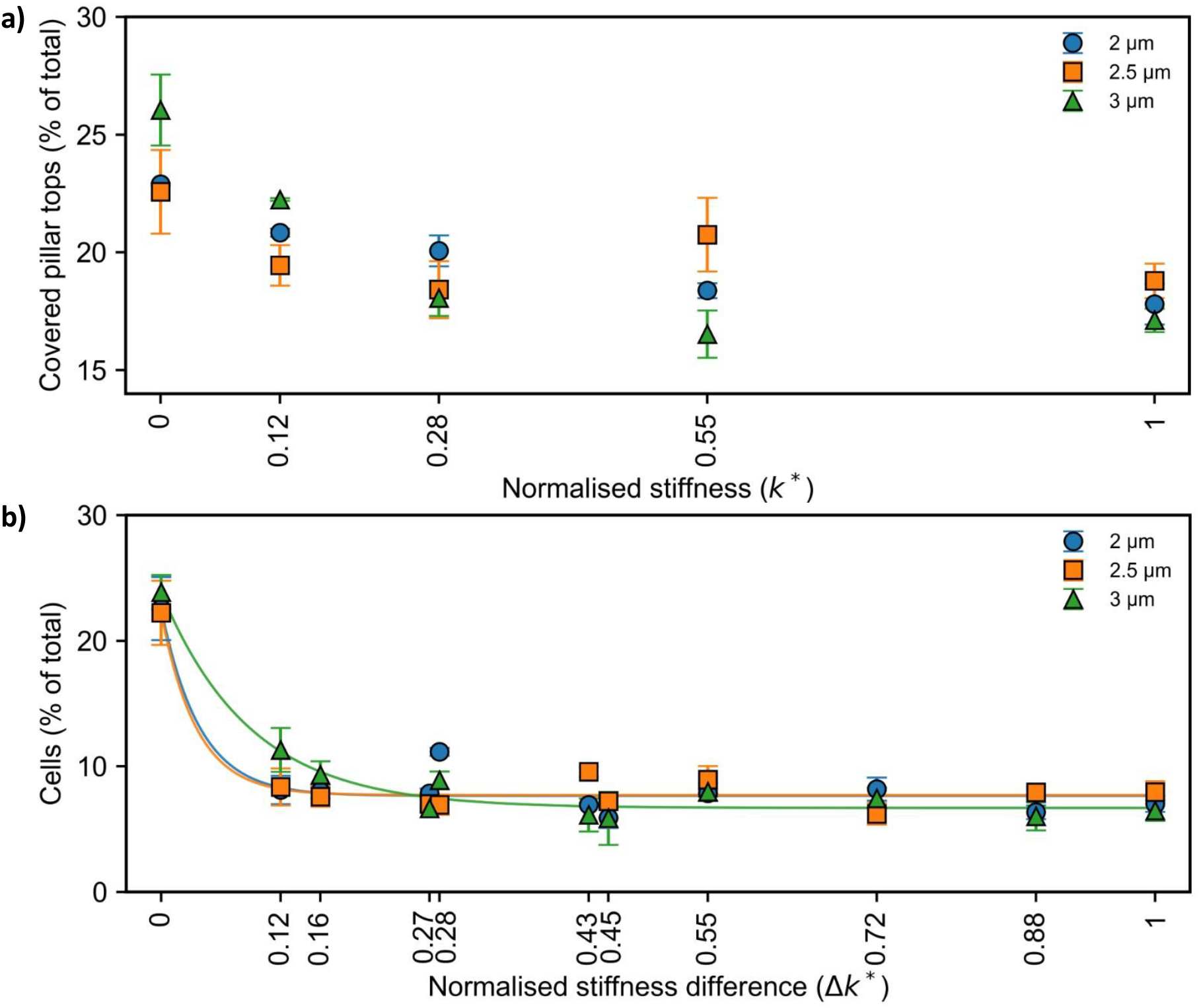
Cell spreading patterns on stiffness-varying micropillar substrates. (a) Distribution of normalised stiffness values (*k*^*\**^) of pillars covered by cells. (b) Distribution of the absolute normalised stiffness difference (Δ*k*^*\**^) between opposite ends of individual cells. Solid lines in (b) represent exponential fits to the corresponding datasets.

To further characterise spreading behaviour, we calculated Δ*k*^*\**^, which represents the absolute normalised stiffness variation experienced across the footprint of a cell. For this analysis, the two ends of an elongated cell were identified as the intersection between the cell major axis and the cell boundary. **Fig. 4b** shows the distribution of cells across different Δ*k*^*\**^ values. This relationship approximately follows an exponential decay, with a greater proportion of cells observed at lower Δ*k*^*\**^ values. However, owing to the repeating stiffness pattern, Δ*k*^*\**^ = 0 can arise through multiple spatial configurations, including cells whose ends occupy the same physical stiffness row as well as cells whose ends occupy different rows with the same stiffness.

To test the distribution expected from substrate geometry alone, a Monte Carlo randomisation with 10,000 iterations was performed separately for each biological replicate. The number of cells and substrate geometry were retained, while cell centroid position and major-axis orientation were randomised, and cell length was sampled within the experimentally observed range for each replicate.

The randomised distribution produced approximately 18% of cells with Δ*k*^*\**^ = 0. However, only ∼14% of these randomly occurring Δ*k*^*\**^ = 0 configurations arose from cells with both ends positioned on the same physical stiffness row, corresponding to alignment along the CSD. In comparison, approximately 23% of experimentally observed cells exhibited Δ*k*^*\**^ = 0, of which 48–50% arose from same-row configurations. These percentage values represent pooled mean values across replicates. Thus, although the overall occurrence of Δ*k*^*\**^ = 0 was only moderately higher than that expected from the substrate geometry alone, its spatial origin differs substantially from the random distribution. The enrichment of same-row configurations indicates that cells preferentially adopt spreading configurations that minimise stiffness variation across their bodies, suggesting that cells tend to avoid large local stiffness differences during spreading.

These observations may be associated with the cell traction-force response discussed earlier. As shown in Section 3.1, cells tend to maintain a relatively consistent level of substrate deformation across a range of stiffness conditions. At the same time, cellular traction forces must remain mechanically balanced, as cells do not sustain a persistent, unbalanced net force. Because cells maintain a comparable level of pillar deformation across different stiffness values, the force on each stiffer pillar is greater than that generated by a softer pillar. Therefore, attachment to a few stiffer pillars may be balanced by attachment to a greater number of softer pillars. This may explain the large number of soft pillars observed to be covered by cells (**Fig. 4a**). In addition, because the relative stiffness gradient increases toward the stiffer region, maintaining a low Δ*k*^*\**^ becomes progressively more favourable when cells spread within the softer regions. This provides further support for the preferential contact with a higher number of softer pillars observed in **Fig. 4a**.

### 3.3 Morphological Response

Cells exhibited distinct pillar-to-pillar spreading behaviour across the pillar array. Notably, cells preferred to elongate along the horizontal and vertical directions rather than along intermediate (diagonal) directions (**Fig. 5a,b**). Orthogonal alignment was quantified separately as the percentage of cells oriented within the 0–10° and 80–90° ranges, together with a combined alignment fraction incorporating both ranges (**Fig. 5c**). This preferential spreading behaviour may arise from a combination of contact guidance, as reported previously [33], and the need to maintain CTF balance across local stiffness gradients. Maintaining a lower Δ*k*^*\**^ favours spreading either along the stiffness-varying direction until regions of similar stiffness are reached or along the constant-stiffness direction, where stiffness variation across the cell footprint remains minimal. In contrast, diagonal spreading generally requires greater elongation to bridge comparable stiffness regions while maintaining force balance. Together, these geometrical and mechanical constraints promote preferential alignment along the two orthogonal directions.

**Fig. 5.**
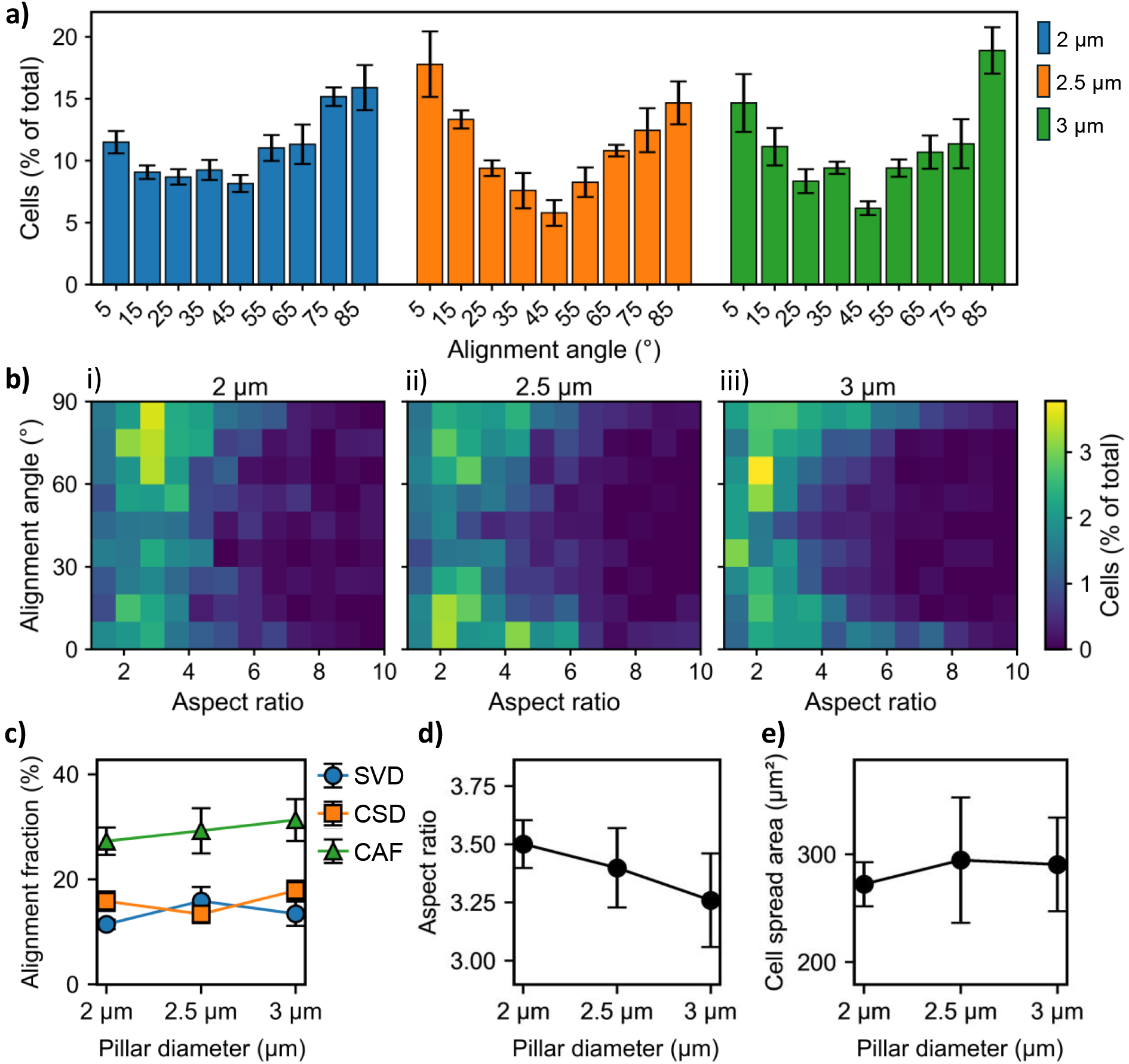
Alignment behaviour on stiffness-varying pillar arrays. (a) Histogram of the cell major-axis orientation angle measured relative to the stiffness-varying direction. An angle of 0° represents cells fully aligned with the stiffness-varying direction, while 90° represents cells aligned with the constant-stiffness direction. (b) Relationship between alignment angle and aspect ratio of the cells. Aspect ratio was calculated as the ratio between the major and minor axes of the fitted ellipse around a cell. (c) SVD, CSD and combined (CAF) alignment fractions as a function of pillar diameter. The SVD and CSD fractions are the percentages of cells oriented within 0–10° and 80–90° of the stiffness-varying direction, respectively, and CAF is their sum. (d) Cell aspect ratio as a function of pillar diameter. (e) Cell spread area variation with pillar diameter. Data points represent pooled values calculated from all analysed cells within each pillar diameter group. Error bars represent the standard error of the mean calculated from three independent experimental replicates (*n* = 3).

Cells on the 2-µm-pillar arrays preferred to elongate along the constant-stiffness direction, whereas cells on the 2.5-µm- and 3-µm-pillar arrays strongly preferred the two orthogonal directions, with no clear preference between them. As shown in **Fig. 3a**, the biphasic displacement response suggests that, on the 2-µm-pillar arrays, the softer pillars may provide insufficient mechanical resistance to support effective focal adhesion maturation. Under these conditions, spreading across regions containing both stiff and soft pillars may be less favourable. The constant-stiffness direction provides a more mechanically homogeneous environment. Here, cells interact with pillars of similar stiffness and do not need to attach to pillars of different stiffness to maintain CTF balance. In contrast, on the 2.5-µm- and 3-µm- pillar arrays, cells exhibit approximately constant substrate deformation across different stiffness values. A lower Δ*k*^*\**^ can be maintained either by spreading along the constant-stiffness direction or the stiffness-varying direction, while remaining in a region of relatively similar stiffness. Consistent with this increased preference for the principal directions of the stiffness pattern, the percentage of orthogonally spread cells, quantified by the combined alignment fraction (CAF), gradually increased from 27.3% to 31.3% with increasing pillar diameter (**Fig. 5c**). The greater stiffness difference between adjacent pillar rows at larger pillar diameters may further reinforce this orthogonal alignment by making intermediate orientations, which expose the cell to larger stiffness variations across its footprint, mechanically less favourable.

**Fig. 5b** shows the relationship between cell alignment angle and aspect ratio for substrates with different pillar diameters. The heatmaps exhibit a characteristic “C”-shaped distribution, where cells with higher aspect ratios prefer to align along the two orthogonal directions. This behaviour becomes more pronounced with increasing pillar diameter. As discussed above, intermediate orientations may become mechanically less favourable as the stiffness difference between adjacent levels increases, potentially restricting highly elongated cells towards the two principal directions of the stiffness pattern. However, the overall cell aspect ratio shows a slight decrease with increasing pillar diameter (**Fig. 5d**), accompanied by a slight increase in cell spread area (**Fig. 5e**). This observation suggests that increasing overall substrate stiffness does not simply promote cell elongation but may also enhance overall cell spreading. Our results are consistent with previous reports of enhanced cell spreading on stiffer substrates [7, 8].

## 4. Conclusion

In this study, we experimentally investigated the behaviour of MDA-MB-231 breast cancer cells on stiffness-varying pillar arrays of different diameters. Varying pillar length modulates the substrate stiffness experienced by cells while maintaining a flat-top contact geometry and constant pillar spacing. Additionally, this platform enables the direct measurement of live-cell traction forces in the presence of steep stiffness gradients within individual cell footprints. Pillar displacement exhibited a biphasic response. In the first phase, displacement decreased gradually at low stiffness. In the second phase, the displacement remained almost constant. In contrast, the calculated CTFs increased across the full stiffness range, with a reduced rate of increase at higher stiffness values. This suggests that cells attempt to reach a mechanoregulated state by maintaining a preferred level of substrate deformation. However, on highly compliant pillars, cells could not maintain this level of deformation, likely due to insufficient substrate resistance to support stable focal adhesion maturation and effective force transmission.

Morphologically, cells remained elongated, although their aspect ratio decreased slightly with increasing pillar diameter. Cells were primarily aligned along the stiffness-varying and constant-stiffness directions. Analysis of cell spreading behaviour showed that cells tend to minimise the difference in stiffness between their two ends and preferentially maintain contact with a larger number of soft pillars. While cells are generally known to prefer stiffer regions, the requirement to maintain a consistent level of substrate deformation and overall force balance may drive cells to engage more with compliant pillars and adopt orthogonal spreading behaviour.

Our findings suggest that cell behaviour on stiffness-varying micropillar arrays is governed by a combination of contact guidance, stiffness sensing, and force-regulation mechanisms. The interplay among these factors leads to distinct spreading patterns and traction-force responses that cannot be explained by stiffness or geometry alone. This work provides insight into how cells respond to microscale stiffness heterogeneity and highlights the importance of considering local force balance in the design of biomimetic substrates for studying cell mechanobiology.

## CRediT authorship contribution statement

Prabuddha De Saram: Methodology, Investigation, Software, Formal analysis, Data curation, Visualisation, Writing – original draft. Nam-Trung Nguyen: Supervision, Resources, Writing – review & editing. Sharda Yadav: Methodology. Navid Kashaninejad: Conceptualisation, Supervision, Funding acquisition, Project administration, Resources, Writing – review & editing.

### Data Availability Statement

The data that support the findings of this study are available from the corresponding author upon reasonable request.

## Appendix A. Supplementary material

Supplementary material associated with this article, including PDMS tensile testing, the tapered-pillar beam model, and the Python script used to calculate pillar stiffness, is available in the online version of this article.

## Acknowledgements

This work used the Queensland node of the NCRIS-enabled Australian National Fabrication Facility (ANFF).

P. De Saram acknowledges Azeez Bakare for his technical assistance during the experimental work.

