## Supplementary Information for "Cellular Traction and Morphological Adaptation to Cell-Scale Stiffness Gradients on Length-Encoded Micropillar Arrays"

**Contents:**

**Section S1. Pillar stiffness calculation**

S1.1 Young's modulus of PDMS (Figs S1–S2)

S1.2 Taper geometry and beam model (Fig. S3 and Table S1)

S1.3 Python Script for micropillar stiffness calculation

### Section S1. Pillar stiffness calculation

#### S1.1 Young's modulus of PDMS

The Young's modulus of the 7:1 PDMS formulation used for micropillar fabrication was experimentally determined by uniaxial tensile testing. An open mould was fabricated from PETG using 3D printing, and uncured PDMS was poured into the mould (**Fig. S1a**). A glass slide was placed over the mould to level the upper surface and produce specimens with approximately uniform thickness. The PDMS was then cured overnight at 60 °C and carefully demoulded after curing.

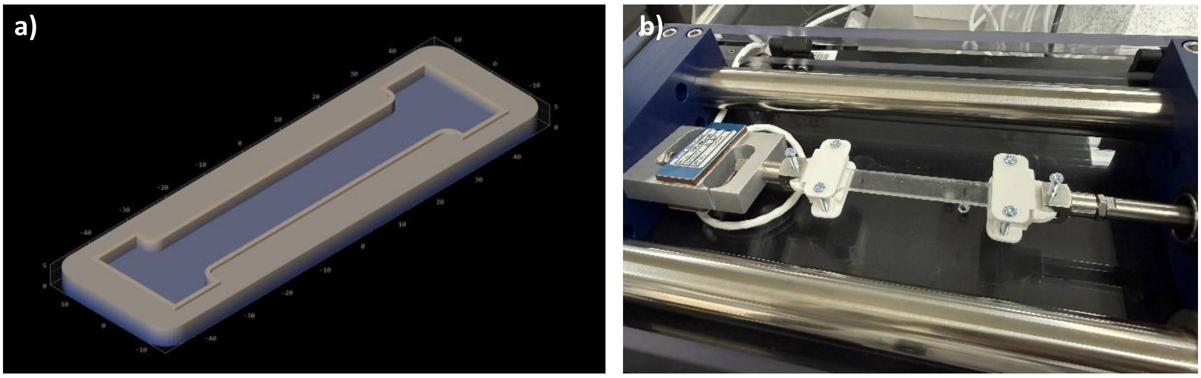


**Fig. S1**. a) Mould design for PDMS specimen. b) Experimental setup

The PDMS specimens were designed with a thickness of 2 mm, a width of 12 mm, and a gauge length of 60 mm. The dimensions of each PDMS specimen were measured using a digital vernier calliper with a resolution of 0.01 mm. The initial cross-sectional area, A, was calculated from the measured specimen width, w, and thickness, t, as $A = wt$.

The specimens were mounted onto the delaminator machine using custom 3D-printed clamps. Each specimen was positioned such that only the uniform cross-sectional region was subjected to stretching. Uniaxial extension was then applied at a constant displacement rate of 50 μm s$^{-1}$ over a total displacement of 1 cm. Force and displacement were recorded throughout the test.

Stress, σ, was calculated from the measured tensile force, F, and the initial specimen cross-sectional area as

$$\sigma = \frac{F}{A} \qquad \text{S1}$$

Strain, ε, was determined from the change in specimen length, ΔL, relative to its initial gauge length, $L_0$, as

$$\varepsilon = \frac{\Delta L}{L_0} \qquad \text{S2}$$

The Young's modulus, E, was determined from the slope of the linear stress–strain relationship,

$$E = \frac{d\sigma}{d\varepsilon} \qquad \text{S3}$$

or, equivalently, from the linear regression

$$\sigma = E\varepsilon + C \tag{S4}$$

where C is the fitted intercept.

Three independent measurements were performed on three different substrates.
**Fig. S2** shows the Stress–strain behaviour.

The calculated Young's modulus values were approximately 2.10, 2.27, and 2.20 MPa. The mean Young's modulus is 2.19 ± 0.08 MPa

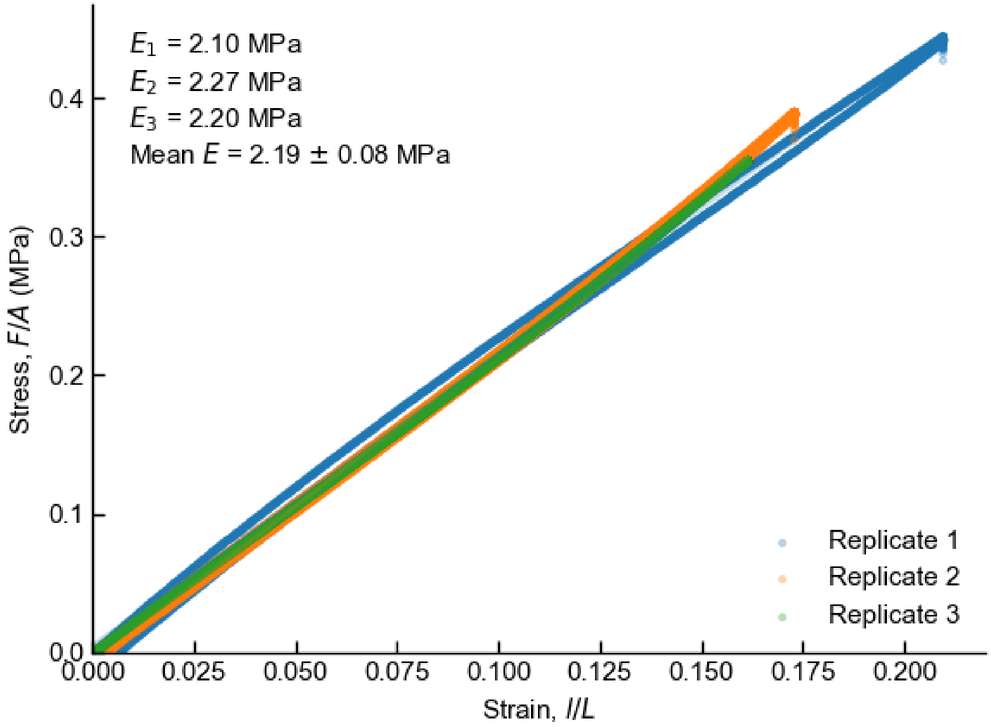


**Fig. S2**. Stress–strain behaviour of 7:1 PDMS measured by uniaxial tensile testing. Three independent measurements are shown together with their corresponding linear regression fits.

This value was approximated as 2.2 MPa for subsequent micropillar stiffness calculations.

#### S1.2 Taper geometry and beam model

SEM images showed that the pillar diameter increased near the base. This tapered region formed during two-photon lithography (2PP) of the moulds. The pillar length matched the design. The pillar geometry used in the stiffness model is shown in **Fig. S3**. For each nominal diameter ($D$) (2.0, 2.5 and 3.0 μm), the following parameters were measured from SEM images and average values were used for the calculations.

- Base diameter ($D_b$)
- Taper angle (θ)

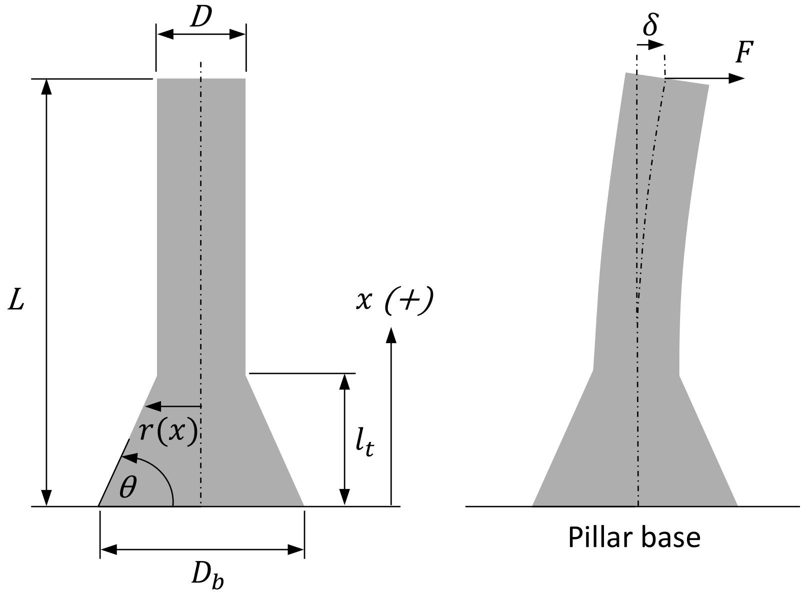


**Fig. S3.** Pillar measurement convention used in the calculations.

**Table S1** summarises the measured dimensions used in the stiffness calculations. For each diameter value, these taper dimensions do not vary significantly with pillar length from base to top (L).

The height of the tapered region ($l_t$),

$$l_t = \frac{D_b - D}{2}\tan\theta \qquad \text{S5}$$

The micropillar local radius is $r(x)$,

$$r(x) = \begin{cases} r_b - x/tan(\theta), & 0 \leq x \leq l_t \\ r, & l_t \leq x \leq L \end{cases} \qquad \text{S6}$$

where $r_b = D_b/2$, $r = D/2$. For example, for $D$ = 2.0 μm ($D_b$ = 4.4 μm, $\theta$ = 51.5°), $l_t$ = 1.5 μm.

The second moment of area then becomes:

$$I(x) = \frac{\pi r(x)^4}{4} \qquad \text{S7}$$

Each pillar was modelled as a cantilever fixed at its base and loaded by a lateral force, $F$, at its top.

Euler–Bernoulli beam theory,

$$\frac{d^2y}{dx^2} = \frac{M(x)}{EI(x)} \qquad \text{S8}$$

Substituting the internal bending moment at a distance $x$, under force F, $M(x) = F(L - x)$,

$$\frac{d^2y}{dx^2} = \frac{F(L - x)}{EI(x)} \qquad \text{S9}$$

Integrating twice gives the lateral displacement of the pillar top, $\delta$,

$$\delta = \frac{F}{E}\int_0^L \frac{(L - x)^2}{I(x)} dx \qquad \text{S10}$$

Stiffness $k$,

$$k = \frac{F}{\delta} = \frac{\pi E}{4\int_0^L \frac{(L - x)^2}{r(x)^4} dx} \qquad \text{S11}$$

Because $r(x)$ varies along the pillar, the integral over the tapered region was evaluated numerically (trapezoidal rule, 20,000 points), and the integral over the uniform cylindrical section was evaluated analytically as $(L - l_t)^3/(3I)$. The resulting stiffness values are listed in **Table S1** and were used in the main text (Table 1).

**Table S1**. Micropillar dimensions and estimated stiffness levels ($E$ = 2.2 MPa).

| D (µm) | $D_b$ (µm) | $\theta$ (°) | $l_t$ (µm) | $L$ (µm) | $k$ (nN/µm) |
|---|---|---|---|---|---|
| 2.0 | 4.4 | 51.5 | 1.5 | 8.0 | 16.0 |
| | | | | 9.0 | 10.6 |
| | | | | 10.0 | 7.4 |
| | | | | 11.0 | 5.4 |
| | | | | 12.0 | 4.0 |
| 2.5 | 5.0 | 49.1 | 1.5 | 8.0 | 37.0 |
| | | | | 9.0 | 24.8 |
| | | | | 10.0 | 17.4 |
| | | | | 11.0 | 12.7 |
| | | | | 12.0 | 9.5 |
| 3.0 | 5.6 | 54.0 | 1.8 | 8.0 | 82.6 |
| | | | | 9.0 | 54.9 |
| | | | | 10.0 | 38.3 |
| | | | | 11.0 | 27.8 |
| | | | | 12.0 | 20.8 |

#### S1.3 Python Script for micropillar stiffness calculation

The script below reproduces Table S1 (Python 3, NumPy ≥ 2.0, pandas).

```
from pathlib import Path
import numpy as np
import pandas as pd

YOUNGS_MODULUS_MPA = 2.2
PILLAR_HEIGHTS_UM = [8.0, 9.0, 10.0, 11.0, 12.0]

PILLAR_GEOMETRIES = {
    2.0: {
        "base_diameter_um": 4.441,
        "taper_angle_deg": 51.52,
    },
    2.5: {
        "base_diameter_um": 5.016,
        "taper_angle_deg": 49.12,
    },
    3.0: {
        "base_diameter_um": 5.627,
        "taper_angle_deg": 53.95,
    },
}

INTEGRATION_POINTS = 20_000
OUTPUT_CSV = Path("micropillar_stiffness_results.csv")

def second_moment_of_area(radius_m):
    return np.pi * np.asarray(radius_m) ** 4 / 4.0

def calculate_taper_height(
    nominal_diameter_um,
    base_diameter_um,
    taper_angle_deg,
):
    if base_diameter_um < nominal_diameter_um:
        raise ValueError("Base diameter cannot be smaller than nominal diameter.")

    if not 0.0 < taper_angle_deg < 90.0:
        raise ValueError("Taper angle must be between 0 and 90 degrees.")

    radial_extension_um = (base_diameter_um - nominal_diameter_um) / 2.0
    return radial_extension_um * np.tan(np.deg2rad(taper_angle_deg))


def calculate_stiffness(
    nominal_diameter_um,
    base_diameter_um,
    taper_angle_deg,
    pillar_height_um,
    youngs_modulus_mpa,
    integration_points,
```

```
):
    taper_height_um = calculate_taper_height(
        nominal_diameter_um,
        base_diameter_um,
        taper_angle_deg,
    )

    if taper_height_um >= pillar_height_um:
        raise ValueError("Calculated taper height must be smaller than pillar height.")

    nominal_radius_m = nominal_diameter_um * 1e-6 / 2.0
    base_radius_m = base_diameter_um * 1e-6 / 2.0
    taper_height_m = taper_height_um * 1e-6
    pillar_height_m = pillar_height_um * 1e-6
    youngs_modulus_pa = youngs_modulus_mpa * 1e6

    x_taper = np.linspace(0.0, taper_height_m, integration_points)
    radius_taper = base_radius_m + (
        nominal_radius_m - base_radius_m
    ) * (x_taper / taper_height_m)

    moment_taper = second_moment_of_area(radius_taper)
    taper_integrand = (pillar_height_m - x_taper) ** 2 / moment_taper
    taper_integral = np.trapezoid(taper_integrand, x_taper)

    nominal_moment = second_moment_of_area(nominal_radius_m)
    cylindrical_integral = (
        pillar_height_m - taper_height_m
    ) ** 3 / (3.0 * nominal_moment)

    total_integral = taper_integral + cylindrical_integral
    stiffness_n_per_m = youngs_modulus_pa / total_integral

    return {
        "nominal_diameter_um": nominal_diameter_um,
        "base_diameter_um": base_diameter_um,
        "taper_angle_deg": taper_angle_deg,
        "taper_height_um": taper_height_um,
        "pillar_height_um": pillar_height_um,
        "youngs_modulus_mpa": youngs_modulus_mpa,
        "stiffness_nN_per_um": stiffness_n_per_m * 1000.0,
    }


def main():
    results = []

    for nominal_diameter_um, geometry in PILLAR_GEOMETRIES.items():
        for pillar_height_um in PILLAR_HEIGHTS_UM:
            results.append(
                calculate_stiffness(
                    nominal_diameter_um=nominal_diameter_um,
                    base_diameter_um=geometry["base_diameter_um"],
                    taper_angle_deg=geometry["taper_angle_deg"],
                    pillar_height_um=pillar_height_um,
                    youngs_modulus_mpa=YOUNGS_MODULUS_MPA,
                    integration_points=INTEGRATION_POINTS,
```

```
            )
        )

    dataframe = pd.DataFrame(results)
    dataframe.to_csv(OUTPUT_CSV, index=False)

    print(dataframe.round(4).to_string(index=False))
    print(f"\nSaved: {OUTPUT_CSV.resolve()}")

if __name__ == "__main__":
    main()
```
